# Multichannel Silicon Probes for Awake Hippocampal Recordings in Large Animals

**DOI:** 10.1101/400812

**Authors:** Alexandra V. Ulyanova, Carlo Cottone, Christopher D. Adam, Kimberly G. Gagnon, D. Kacy Cullen, Tahl Holtzmann, Brian G. Jamieson, Paul F. Koch, H. Isaac Chen, Victoria E. Johnson, John A. Wolf

## Abstract

Decoding laminar information across deep brain structures and cortical regions is necessary in order to understand the neuronal ensembles that represent cognition and memory. Large animal models are essential for translational research due to their gyrencephalic neuroanatomy and significant white matter composition. A lack of long-length probes with appropriate stiffness to penetrate to deeper structures with minimal damage to the neural interface is one of the major technical limitations to applying the approaches currently utilized in lower order animals to large animals. We therefore tested the performance of multichannel silicon probes of various solutions and designs that were developed specifically for large animal electrophysiology. Neurophysiological signals from dorsal hippocampus were recorded in chronically implanted awake behaving Yucatan pigs. Single units and local field potentials were analyzed to evaluate performance of given silicon probes over time. EDGE-style probes had the highest yields during intra-hippocampal recordings in pigs, making them the most suitable for chronic implantations and awake behavioral experimentation. In addition, the cross-sectional area of silicon probes was found to be a crucial determinant of silicon probe performance over time, potentially due to reduction of damage to the neural interface. Novel 64-channel EDGE-style probes tested acutely produced an optimal single unit separation and a denser sampling of the laminar structure, identifying these research silicon probes as potential candidates for chronic implantations. This study provides an analysis of multichannel silicon probes designed for large animal electrophysiology of deep laminar brain structures, and suggests that current designs are reaching the physical thresholds necessary for long-term (~ 1 month) recordings with single-unit resolution.

## Introduction

Spatiotemporal neuronal ensembles distributed throughout laminar structures in the brain such as the cortex and hippocampus are presumed to be the substrate for cognition and memory. Oscillatory activity and resulting power bands are transmitted throughout the layers of laminar structures as different inputs are driven at different frequencies depending on their area of origin [1–5]. Laminar analyses of oscillatory activity have proven valuable for elucidating circuit dynamics as well as changes between states in both cortex and hippocampus [5, 6]. Traditionally, single neurons have been analyzed independently on the basis of their tuning to sensory stimuli or movement. Although accuracy of tuning curve approaches is unaffected by growing numbers of simultaneously recorded neurons, newly developed pair-wise interaction models that make predictions based on the activity of the multiple simultaneously recorded neurons become more complex but also more accurate as the number of recorded neurons increases [7]. For spike-field entrainment analyses as well as for neuromodulation-based approaches, it is also important to know where inputs that drive the cells originate and how they interact locally. Cell types, dendritic arborization, local and long-range axonal projections are all distributed unevenly across layers, shaping the integration and segregation of neural signals. Ultimately, we must decode laminar information across different structures to understand the spatiotemporal ensembles that represent cognition and memory. Thus, measuring the coordination of spiking activity of large numbers of neurons, and specifically those thought to give rise to distributed functional networks, is critical for understanding neural information processing underlying cognition and behavior [6].

The hippocampus is an example of a deep laminar structure that is highly involved in encoding episodic memory formation presumably via spatiotemporal ensembles of neurons [8, 9]. Once multichannel silicon probes were developed, they quickly became a standard tool for awake hippocampal neurophysiological recordings in rodents [10]. Increasing the density of laminar contacts in deep structures allows researchers to gain more information about these structures and more recently, study the potential effects of neurological disease states on single unit activity [11–13]. Although the rodent hippocampus has been studied in great detail, the transition to large animal models has been slower. One of the major obstacles to addressing the research questions and approaches currently accessible in rodents are technical limitations in laminar silicon microelectrodes. While translational models have been actively utilized for neuromodulation studies [14–18], much of the large animal brain is still inaccessible due to insufficient length of available silicon probes. In large animals and humans, the hippocampus is located sub-cortically and therefore difficult to study electrophysiologically using multi-channel probes. Even many cortical targets cannot be reached with the current technology including those typically utilized for brain machine interface (BMIs) such at the Utah electrode. In addition, laminar structure cannot be detected with single wire or tetrode recordings. Existing non-laminar solutions can scale to high density, but have single contacts at the tip, limiting the ability to differentiate single units or localize them within the laminar circuitry [19]. Therefore, current technology is not suitable for understanding neuronal oscillation and their interaction within and between laminar structures.

Recent advances in high-density linear electrode arrays and wireless recording technology could tremendously enhance translational research studies of large-scale networks in species with large, gyrencephalic brains [20]. Ideally, the probes for electrophysiological recordings from deep brain structures must resolve laminar local field potentials (LFPs) and provide proper sampling densities to resolve single-units across channels for single unit spike-sorting and minimize damage to the neuro-electric interface (for review see [21]). For instance, multi-channel silicon probes with many contacts in a linear configuration can isolate units in a 2 – 300 μm “sphere” around each electrode [22, 23]. Layer-specific fields can convey information transmitted from other regions, while direct, local spiking activity can be measured simultaneously. Silicon probes also need to be long enough to reach these structures. With little damage to the neural interface, insertion of silicon into multiple brain regions probes could also provide highly detailed information about local and long-range circuit computations [6].

Technology necessary to produce finer features at longer length has only recently became available, including stitching across the dye in the photomask process to utilize smaller feature processes. New multichannel silicon probes designed for large animal electrophysiology allow for simultaneous recordings of fields and spikes from multiple layers of laminar structures such as cortex and hippocampus, and we therefore tested various designs of laminar silicon probes suitable for large animals. Across probe designs, we compared neurophysiological characteristics and biomechanical compatibility based on the variables such as electrode site layout and probe thickness. Current technology in silicon probes appears to have reached the critical size/feature dimension while maintaining insertion stiffness, proving that single units in deep brain structures can be recorded for chronic periods without extensive damage to the neural interface. Further refinement of these probe technologies in combination with chronic drives and wireless technology may usher in a new high-density era for previously unexplored regions of cortex, hippocampus and other non-laminar deep structures in large animals and potentially humans.

## Materials and Methods

### Animals

Male Yucatan miniature pigs were purchased from Sinclair (NSRRC, Catalog # 0012, RRID: NSRRC_0012) and underwent the current studies at the approximate age of 5 – 6 months at a mean weight of 38 ± 3 kg (n = 17, mean ± SEM). At this age Yucatan pigs are considered post-adolescent with near-fully developed brains, while young enough to be of a manageable weight for procedures and behavior [24–26]. All pigs were pair housed when possible, and were always in a shared room with other pigs. All animal procedures were performed in accordance with the University of Pennsylvania animal care committee’s regulations, an AALAC accredited institution.

### Surgical Procedure

Yucatan miniature pigs were fasted for 16 hours then induced with 20 mg/kg of ketamine (Catalog # NDC 0143-9509-01, West-Ward, Eatontown, NJ) and 0.5 mg/kg of midazolam (Catalog # NDC 0641-6060-01, West-Ward, Eatontown, NJ). Animals were intubated with an endotracheal tube and anesthesia was maintained with 2 – 2.5 % isoflurane per 2 liters O2. Each animal was placed on a ventilator and supplied oxygen at a tidal volume of 10 mL/kg. A catheter was placed in an auricular vein to deliver 0.9 % normal saline at 200 mL per hour. Additionally, heart rate, respiratory rate, arterial oxygen saturation, end tidal CO_2_ and rectal temperature were continuously monitored, while pain response to pinch was periodically assessed. All of these measures were used to titrate ventilation settings and isoflurane percentage to maintain an adequate level of anesthesia. A forced air warming system was used to maintain normothermia throughout the procedure.

All animals underwent implantation of multi-channel silicon probe under sterile conditions similar to acute implantation procedure described previously [20]. Briefly, pigs were placed in a stereotactic frame, with the surgical field prepped and draped, and a linear incision was made along the midline. A 13-mm diameter burr hole was centered at 7 mm lateral to the midline and 4.5 mm posterior to bregma, and the bony opening was subsequently expanded using Kerrison punches. Skull screws were placed over the occipital ipsilateral and contralateral cortex as a ground and alternate reference signal. The dura was opened in a cruciate manner and the brain was mapped in the sagittal plane with a tungsten electrode (impedance = 0.5 MΩ, measured at 1KHz; Catalog # UEWSEGSEBNNM), utilizing observed spiking activity to generate a two-dimensional map. Based on this map, a silicon probe was inserted so that the spread of electrode sites would span the laminar structure of the dorsal hippocampus at its maximal thickness in the dorsal-ventral plane, perpendicular to the individual hippocampal layers [20]. The multichannel silicon probe was stabilized inside the craniectomy using Tisseel (Catalog # 1504514, Baxter Healthcare, Wayne, PA), creating a semi-floating interface between the silicon probe and the animal’s skull. The probe cables were fed through the head cap chamber and attached to a custom electrode interface board (EIB) adapter. The head cap chamber, which covered all necessary electronic connections, was attached to the pig skull using anchor screws and secured with bone cement and geristore.

Once the quality of signals from multichannel silicon depth probes were confirmed, hippocampal signals were first recorded under anesthesia using a QC-72 amplifier and referenced to both internal and skull references. This recording was later used as a baseline for awake recordings and to monitor silicon probe’s position over time for possible drift.

### Multichannel Silicon Probes

Chronic 32-channel silicon probes for large animal electrophysiology were designed with ATLAS Neuroengineering (Leuven, Belgium) and NeuroNexus (Ann Arbor, MI). In addition, acute 64-channel silicon research probes were developed in a collaboration with SB Microsystems (Glen Burnie, MD) and Cambridge NeuroTech (Cambridge, UK). Designs of multichannel silicon probes are shown in Figure 1. Specific parameters of silicon probes are summarized in Table 1.

**Figure 1.**
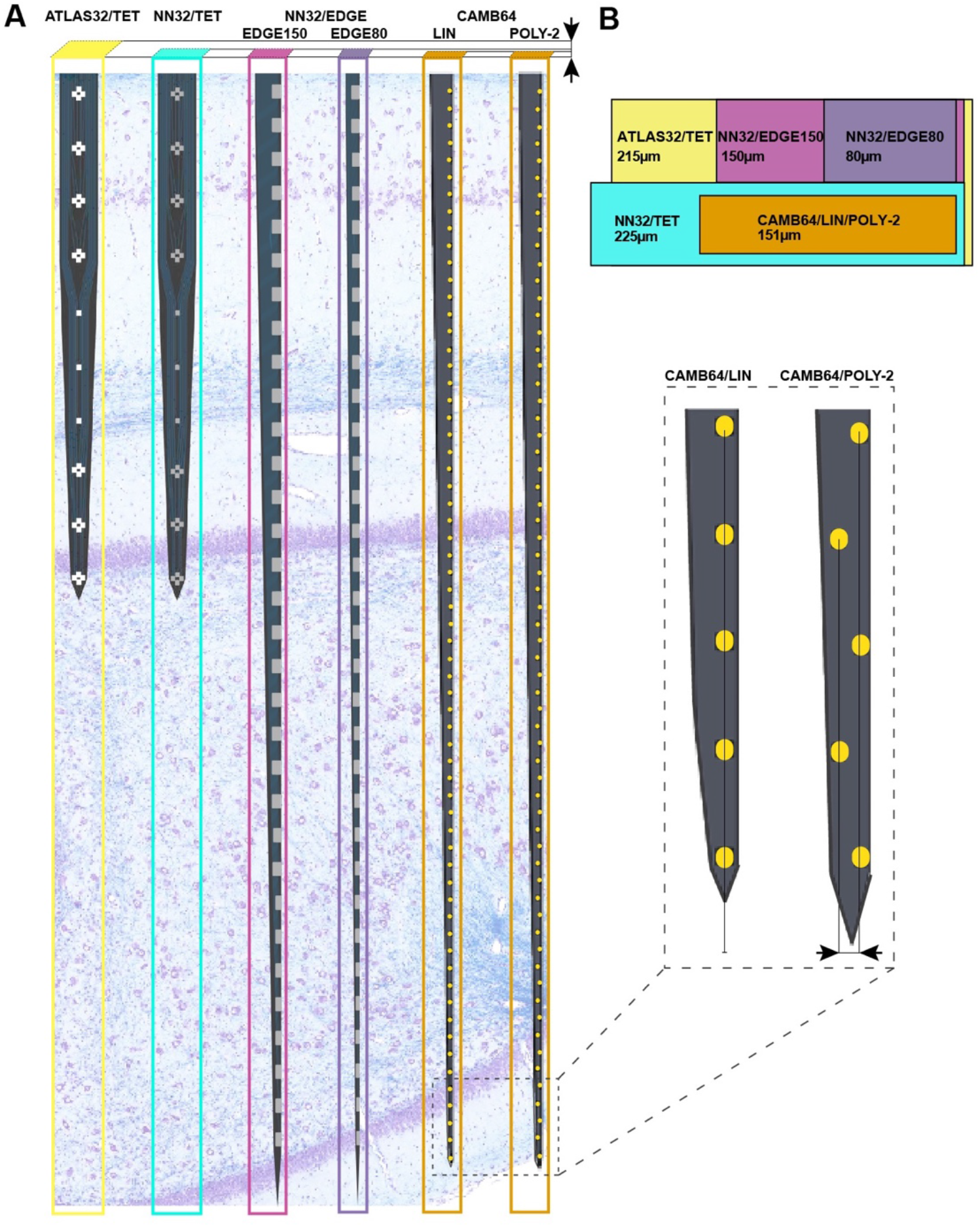
Design of multichannel silicon probes for chronic hippocampal recordings in pigs. **A)** The arrangement of the individual electrode sites is shown for chronic 32-channel silicon probes (ATLAS32/TET (yellow), NN32/TET (blue), NN32/EDGE150 (pink), and NN32/EDGE80 (purple)) and acute 64-channel silicon probes (CAMB64/EDGE and CAMB64/POLY-2, orange). The vertical offset for CAM64 silicon probes is also shown magnified (inset). Silicon probes are shown overlaid on a representative sagittal section of dorsal hippocampus stained with LFB/CV to identify individual layers. Cross-sectional area of each silicon probe (measured at the top electrode site) is shown at the top. **B)** Widths and thicknesses of multichannel silicon probes (also referred to as Probe Area) are shown overlaid on each other for comparison (colors are the same as in A). The corresponding widths are 215 μm for ATLAS32/TET (yellow), 150 μm for NN32/EDGE150 (pink), 80 μm for NN32/EDGE80 (purple), 225 μm for NN32/TET (blue), and 151 μm for CAMB64/EDGE/POLY-2 (orange) silicon probes. NN32/EDGE150 is shown as a continuation of NN32/EDGE80. The thickness of silicon probes is 100 μm for ATLAS32/TET, 50 μm for NN32/EDGE150, NN32/EDGE80, NN32/TET, and 35 μm for both CAMB64/EDGE and CAMB64/POLY-2 probes (orange).

**Table 1.** Multi-channel silicon probes for chronic hippocampal recordings in pigs.

| Silicon Probe | Site Layout | Site Spacing ($\mu\text{m}$) | Site Coating | Site Area ($\mu\text{m}^2$) | Site Imp ($\text{M}\Omega$) | Probe Thickness ($\mu\text{m}$) | Probe Width ($\mu\text{m}$) | Probe Area ($\mu\text{m}^2$) | Probe Length ($\mu\text{m}$) |
| --- | --- | --- | --- | --- | --- | --- | --- | --- | --- |
| **ATLAS32/**<br>**TET** | tetrode<sup>1</sup><br>+ linear | 275<sup>2</sup> | Pt | 962 | 1.29 | 100 | 215 | 21,500 | 2,750 |
| **NN32/**<br>**TET** | tetrode<sup>1</sup><br>+ linear | 275<sup>2</sup> | IrOx | 312 | 1.68 | 50 | 225 | 11,250 | 11,250 |
| **NN32/**<br>**EDGE150** | edge<sup>3</sup> | 200 | IrOx | 312 | 1.81 | 50 | 150 | 7,500 | 7,500 |
| **NN32/**<br>**EDGE80** | edge<sup>3</sup> | 200 | IrOx | 312 | 1.81 | 50 | 80 | 4,000 | 4,000 |
| **CAMB64/**<br>**EDGE** | edge<sup>3</sup> | 100 | Au +<br>polymer | 165 | 0.06 | 35 | 148 | 5,180 | 5,180 |
| **CAMB64/**<br>**POLY-2** | poly-2<sup>4</sup> | 100 | Au +<br>polymer | 165 | 0.06 | 35 | 154 | 5,390 | 5,390 |

^1^ an arrangement of four electrode sites placed close together;

^2^ spacing between individual tetrodes (at first site) as well as between linear sites;

^3^ similar to the linear layout, but electrode sites are positioned at the edge of the substrate;

^4^ similar to the linear layout, but electrode sites are off-set by 21 μm relative to each other.

All multichannel silicon probes were designed with one low-impedance channel placed 1-2 mm above adjacent proximal channel, which was recorded for use as an internal reference (Figure S1). For dorsal hippocampal targeting in pigs, this design provides 31 (or 63) channels for intra-hippocampal recordings with the reference channel being positioned within the temporal horn of the lateral ventricle dorsal to the hippocampus. Detailed information on multichannel silicon probes used in the study is as follows:

**ATLAS32/TET** (Catalog # E32T7-R-275-S01-L25): 32-channel silicon probes had a custom design individual sites arranged in groups of four closely spaced sites or tetrodes (Figure 1A). Three electrode sites were added in between groups of tetrodes to cover porcine hippocampal layers of strata radiatum, lacunosum-moleculare and moleculare. The top four tetrodes were designed to be positioned in pyramidal CA1 layer, while the bottom three tetrodes positioned in granular cell layer. The top site on each tetrode, along with linear sites were placed 275 μm apart formed 10 equally spaced sites for laminar hippocampal recordings. The total coverage of ATLAS32/TET silicon probe was set to 2,750 μm. The cross-sectional area of the probe, measured at the top site was 21,500 μm^2^ (Figure 1B). The individual electrode sites of 962 μm^2^ were coated with Pt [27]. The average site impedance, measured at 1 KHz was 1.29 ± 0.21 MΩ (mean ± SEM, n_sites_ = 155).

**NN32/TET** (Custom design, Catalog # V1x32-80mm-275-tet-177-HP32): 32-channel silicon probes had a custom design individual sites arranged in groups of four closely spaced sites or tetrodes (Figure 1A). Three electrode sites were added in between groups of tetrodes to cover porcine hippocampal layers of strata radiatum, lacunosum-moleculare and moleculare. The top four tetrodes were designed to be positioned in pyramidal CA1 layer, while the bottom three tetrodes positioned in granular cell layer. The top site on each tetrode, along with linear sites were placed 275 μm apart formed 10 equally spaced sites for laminar hippocampal recordings. The total coverage of NN3232/TET silicon probe was set to 2,750 μm. The cross-sectional area of the probe, measured at the top site was 11,250 μm^2^ (Figure 1B). The individual electrode sites of 312 μm^2^ were coated with IrOx. The average site impedance, measured at 1 KHz was 1.68 ± 0.20 MΩ (mean ± SEM, n_sites_ = 403).

**NN32/EDGE150 / NN32/EDGE80** (Catalog # V1x32-Edge-10mm-200-312-Ref): 32-channel silicon probes had a linear site layout, with the electrode sites placed 200 μm apart. The individual electrode sites were also strategically positioned at the edge of the probe (Figure 1A). The total coverage of NN32/EDGE150 and NN32/EDGE80 probes was set to 6,200 μm. The cross-sectional area of the probe, measured at the top site was 7,500 μm^2^ for NN32/EDGE150 and 4,000 μm^2^ for NN32/EDGE80 (Figure 1B). The individual electrode sites of 312 μm^2^ were coated with IrOx. The average site impedance, measured at 1 KHz was 1.68 ± 0.20 MΩ (mean ± SEM, n_sites_ = 403).

**CAMB64/EDGE**: 64-channel silicon probes had a linear site arrangement, with the individual electrode site positioned at the edge of the silicon probe (Figure 1A). The electrode sites were placed 100 μm apart (Figure 1A, inset). The total coverage of CAMB64/EDGE silicon probe was set to 6,300 μm. The cross-sectional area of the probe, measured at the top site was 5,180 μm^2^. The individual electrode site was 165 μm^2^ (Figure 1B). The individual electrode sites were coated with Au and a conducting organic polymer. The average site impedance, measured at 1 KHz was 0.063 ± 0.001 MΩ (mean ± SEM, n_sites_ = 63).

**CAMB64/POLY-2**: 64-channel silicon probes had individual electrode sites arranged in poly-2 style (Figure 1A). The individual electrode sites were placed 100 μm apart, with a 21 μm off-set (Figure 1A, inset). The electrode sites were placed 100 μm apart. The total coverage of CAMB64/POLY-2 silicon probe was set to 6,300 μm. The cross-sectional area of the probe, measured at the top site was 5,390 μm^2^. The individual electrode site was 165 μm^2^ (Figure 1B). The individual electrode sites were coated with Au and a conducting organic polymer. The average site impedance, measured at 1 KHz was 0.064 ± 0.001 MΩ (mean ± SEM, n_sites_ = 63).

### Neural Data Collection and Analysis

Electrophysiological recordings with 32-channel silicon probes (ATLAS32/TET, NN32/TET, NN32/EDGE150, and NN32/EDGE80) were performed in awake behaving animals at various time points at up to 6 months post implantation. Wide bandwidth neural signals were acquired continuously, sampled at 30 kHz with FreeLynx digital acquisition system, amplified and either wirelessly transmitted to Digital Lynx 4SX acquisition system with Cheetah recording and acquisition software during behavioral space recordings or stored to an on-board microSD memory card during home cage recordings (Neuralynx, Inc., Bozeman, MT). Electrophysiological recordings with 64-channel silicon probes (CAMB64/EDGE and CAMB64/POLY-2) were performed acutely under isoflurane anesthesia, with wide band neural signals continuously acquired, sampled at 32 kHz and amplified with Digital Lynx 4SX acquisition system with Cheetah recording and acquisition software (Neuralynx, Inc., Bozeman, MT).

#### Spike Detection and Analysis

Neural signals acquired from the 31 or 63 channels on the silicon probes were bandpass filtered (0.1 Hz to 9 kHz) in real time prior to sampling. Offline spike detection and sorting was performed on the wideband signals using the Klusta package (http://klusta-team.github.io/klustakwik/, RRID: SCR_014480), which was developed for higher density electrodes, and manually refined with KlustaViewa (https://github.com/klusta-team/klustaviewa), or phy (https://github.com/kwikteam/phy) software packages. The Klusta packages are designed to construct putative clusters from all probe channels simultaneously by taking advantage of spike timing and the special arrangement of electrode sites [28]. After manual refinement, resulting single-unit clusters were then imported into Matlab software, version R2017a for visualization and further analysis using custom and built-in routines (<u>MATLAB</u>, RRID:SCR_001622).

#### Analysis of Local Field Potentials (LFPs)

Acquired wideband LFPs recorded from all channels of the silicon probe were down-sampled to 2 kHz for further analysis. Signals were imported into Matlab software, version R2017a (<u>MATLAB</u>, RRID:SCR_001622) and processed using a combination of custom and modified scripts from the freely available Matlab packages FMAToolbox (<u>FMAToolbox, RRID:SCR_015533</u>), Chronux (<u>Chronux</u>, RRID:SCR_005547), and EEGLAB (<u>EEGLAB</u>, RRID:SCR_007292) [29, 30].

### Tissue Handling and Histological Examinations

Histological analyses were performed to identify electrode tracks on brain tissue from male Yucatan miniature pigs. At the study endpoint, transcardial perfusion was performed under anesthesia using 0.9% heparinized saline followed by 10 % neutral buffered formalin (NBF). After further post-fixation for 7 days in 10 % NBF at 4 ° C, the brain was dissected into 5 mm blocks in the coronal plane and processed to paraffin using standard techniques [31, 32]. 8 μm sections were obtained at the level of the hippocampus in and standard hematoxylin & eosin (H & E) staining was performed on all animals to identify electrode tracks. The following additional stains were performed:

#### Luxol Fast Blue / Cresyl Violet (LFB/CV) Staining

Tissue sections were dewaxed in xylenes and rehydrated to water via graded ethanols before being immersed in 1 % LFB solution (Sigma, S3382) at 60 °C for 4 hours. Excess stain was then removed by immersion of sections in 95 % ethanol. Differentiation was performed via immersion in 0.035 % lithium carbonate for 10 seconds followed by multiple immersions in 70 % ethanol until the gray and white matter could be clearly distinguished. Slides were rinsed and counterstained via immersion in preheated 0.1 % CV solution (Sigma, C5042) for 5 minutes at 60 °C. After further rinsing, slides were differentiated in 95 % ethanol with 0.001 % acetic acid, followed by dehydration, clearing in xylenes and cover slipping using cytoseal-60.

#### Van Gieson’s Staining

Tissue sections were dewaxed in xylenes and rehydrated to water via graded ethanols before being immersed in Weigert’s Working Hematoxylin solution, prepared by mixing equal parts of Weigert’s Iron Hematoxylin A (EMS, Catalog # 26044-05) and Weigert’s Iron Hematoxylin B (EMS, Catalog # 26044-15) for 10 minutes. After rinsing in distilled water, tissue sections were stained for 3 minutes in Van Gieson’s solution (EMS, Catalog # 26046-05). After further rinsing, slides were differentiated in 95 % ethanol with 0.001 % acetic acid, dehydrated, cleared in xylenes and coverlipped.

#### Immunohistochemistry (IHC)

IHC labeling was performed according to previously published protocols [31–33]. Briefly, tissue sections were dewaxed and rehydrated as above, followed by immersion in 3 % aqueous hydrogen peroxide for 15 minutes to quench endogenous peroxidase activity. Antigen retrieval was achieved via microwave pressure cooker at high power for 8 minutes, submerged in Tris EDTA buffer (pH 8.0). Sections were then incubated overnight at 4 ° C using antibodies specific for the N-terminal amino acids 66 – 81 of the amyloid precursor protein (APP) (Millipore, Burlington MA, clone 22C11 at 1:80K), GFAP (Leica, Biosystems, Buffalo Grove, IL, GA5 clone at 1:10K, and IBA1 (Wako Chemicals USA Inc. Richmond, VA at 1:7K). Slides were then rinsed and incubated in the relevant species-specific biotinylated universal secondary antibody for 30 minutes at room temperature. Next, application of the avidin biotin complex (Vector Laboratories, Catalog # PK-6200, Lot # RRID: AB_2336826) was performed for 30 minutes, also at room temperature. Lastly, the 3, 3’-diaminobenzidine (DAB) peroxidase substrate kit (Vector Laboratories, Catalog # SK-4100, Lot # RRID: AB_233638) was applied according to manufacturer’s instructions. All sections were counterstained with hematoxylin, dehydrated in graded ethanols, cleared in xylenes, and cover slipped.

### Statistical Analysis

The data was analyzed using Graphpad Prism software, version 7 (<u>GraphPad Prism</u>, RRID:SCR_002798). Single unit waveforms recorded with silicon laminar probes are displayed as mean ± SD. Amplitudes of single units are displayed as mean ± SEM. Impedance of silicon probes is shown as mean ± SEM.

## Results

### Multichannel Silicon Probes Designed for Chronic Implantation in Large Animal

Multichannel silicon probes designed for large animal electrophysiology were evaluated for their ability to continuously record neurophysiological signals (laminar oscillatory and single unit activity) in awake behaving pigs, and for neuropathological changes induced by their placement in the porcine hippocampus over time.

#### Awake Hippocampal Recordings in Large Animals

Using silicon probes of various designs, the laminar structure of pig dorsal hippocampus was examined electrophysiologically under awake behaving conditions (Figure 2). Since the hippocampal region of interest (at our standard recording coordinates in medial-lateral (ML) and anterior-posterior (AP) planes) is about 1,600 μm (see [20] for more details), silicon probes with 200 μm site spacing provide good coverage for the laminar structure with enough resolution to identify most layers (Figure 1A). Oscillatory activity of the porcine hippocampus recorded with NN32/EDGE80 silicon probe is shown in Figure 2 (Site Spacing = 200 μm, Probe Length = 6,200 μm). Neurophysiological features of awake porcine hippocampus such as sharp-wave ripple (SPW-R) and the polarity inversion of the LFPs across stratum radiatum are shown (Figure 2). While this silicon probe allowed us to examine the porcine hippocampus in its entirety (including the deeper layers such as hilus), some of the laminar layers (such as L-M) could have easily been poorly sampled using 200 μm resolution [20].

**Figure 2.**
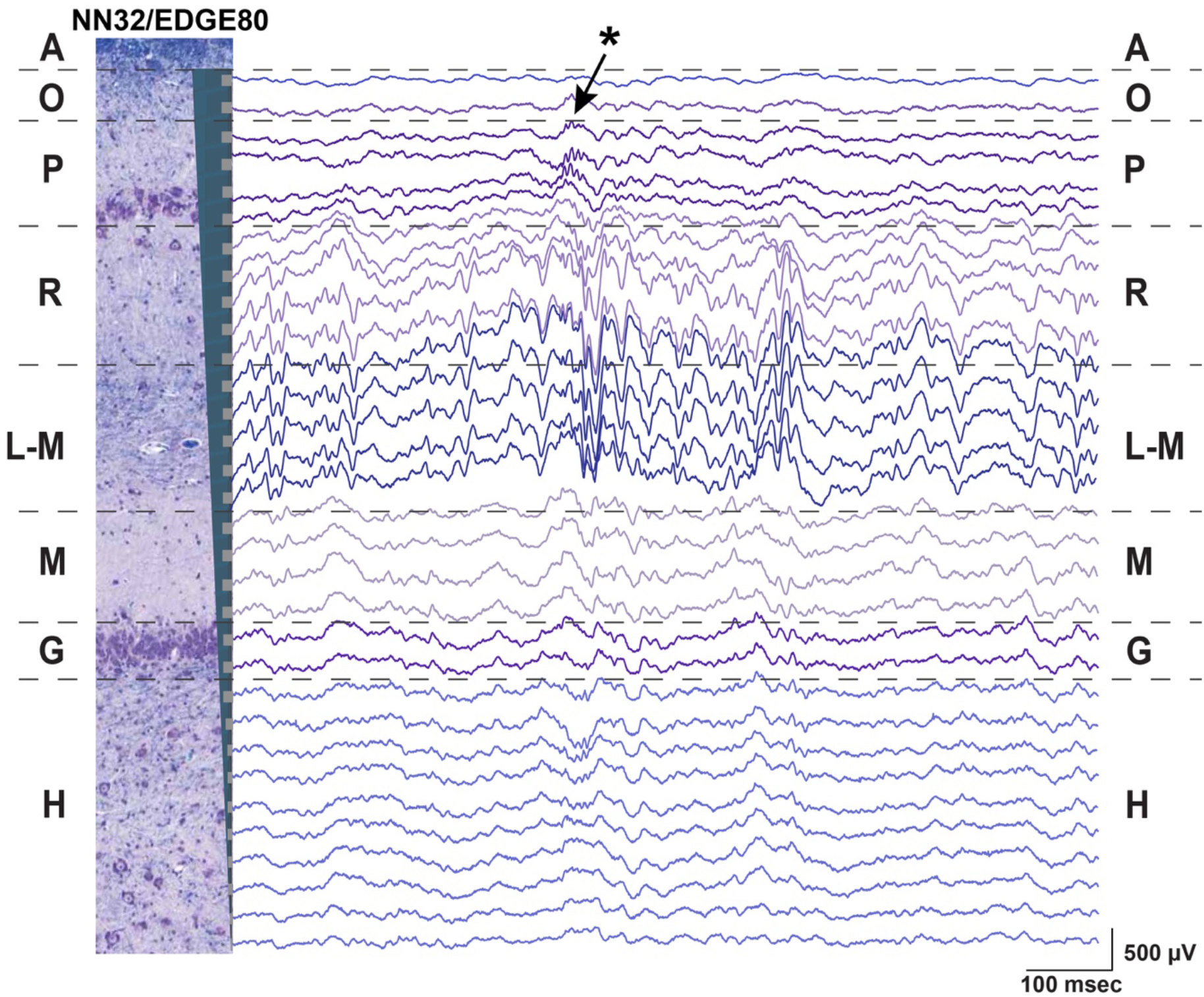
Chronic hippocampal recordings in awake behaving pigs. Hippocampal recording with NN32/EDGE80 silicon probe (Site Spacing = 200 μm) is shown for awake behaving animal. The silicon probe was placed in the dorsal hippocampus so that it covers hippocampal layers of stratum oriens (O), pyramidale (P), radiatum (R), lacunosum-moleculare (L-M), moleculare (M), granulosum (G), and hilus (H). A representative hippocampal section stained with LFB/CV is overlaid on laminar oscillatory activity to match representative layers as described previously [20]. A single sharp-wave ripple event (SPW-R) can be seen in CA1 layer (arrow). The polarity inversion of LFPs across stratum radiatum hippocampal layer can also be seen.

Neurophysiological signals recorded from deep brain structures such as hippocampus may confer noise from the electrical environment by using an electrically inactive but conductive structure such as the cerebrospinal fluid (CSF) of the ventricle. Designing silicon probes with an internal reference for intra-hippocampal (or other deep brain structures) recordings may help to reduce noise and improve the signal quality over time. We estimated how an introduction of the internal reference affects the noise during awake and anesthetized hippocampal recordings by comparing power of hippocampal oscillation at the level of pyramidal CA1 layer referenced to either an internal or skull screw references (Figure S1). All multichannel silicon probes used in the study were custom-designed to have a top electrode site substituted for a low-impedance reference site (Site Area = 4,200 μm^2^) and placed 1-2 mm above the most-proximal probe site (depending on the silicon probe design) (Figure S1, A). For dorsal hippocampal targeting in pigs, these designs provide 31 channels for laminar recordings and results in the reference channel being positioned within the temporal horn of the lateral ventricle sitting just above the hippocampus. During awake behaving recordings in pigs, the internal reference eliminated noise associated with movement artifacts (Figure S1, B and C). Under anesthetized preparation, the internal reference on silicon probes eliminated most of the slow “drift” oscillations as well as 60 Hz frequencies peak, presumably from AC noise during acute recordings, with tether used to record electrophysiological signals (Figure S1, D). The skull screw reference used for large animal awake recordings allows for comparative analysis with a rodent awake behaving literature. In addition, an internal reference may also be beneficial for chronic recordings if a skull screw loses a connection to the CSF due to a growth of the animal’s skull over time (months).

#### Stability of Neuronal Oscillations over Time

We evaluated the stability of neurophysiological signals recorded from pig hippocampus over time in a sub-sample of these probes. Changes in hippocampal signals were evaluated by using power of LFP signals over time (Figure 3). Since theta (~ 4 – 10 Hz in pigs) is considered to be a prominent oscillation in pyramidal CA1 layer of hippocampus, we first calculated power of theta (at the stratum radiatum) over the first two weeks post implantation with NN32/EDGE80 silicon probe (Figure 3A). In the first two weeks following surgery, chronically implanted silicon probes moved slightly, likely due to recovery effects post-surgery such as swelling (Figure 3A). The peak of theta power shifted from channel 8 to channel 5, indicating that NN32/EDGE80 silicon probe moved about 600 μm into the hippocampus, corresponding to a distance between 3 channels on NN32/EDGE80 probe with 200 μm site spacing (Figure 3B). Overall, all previously characterized hippocampal oscillations (low and high gamma, ripple oscillations) decreased over months post-surgery, with the largest drop seen in the first month, potentially due to gliosis/scar tissue formation leading to an insulating/filtering effect on the signal (Figure 3C).

**Figure 3.**
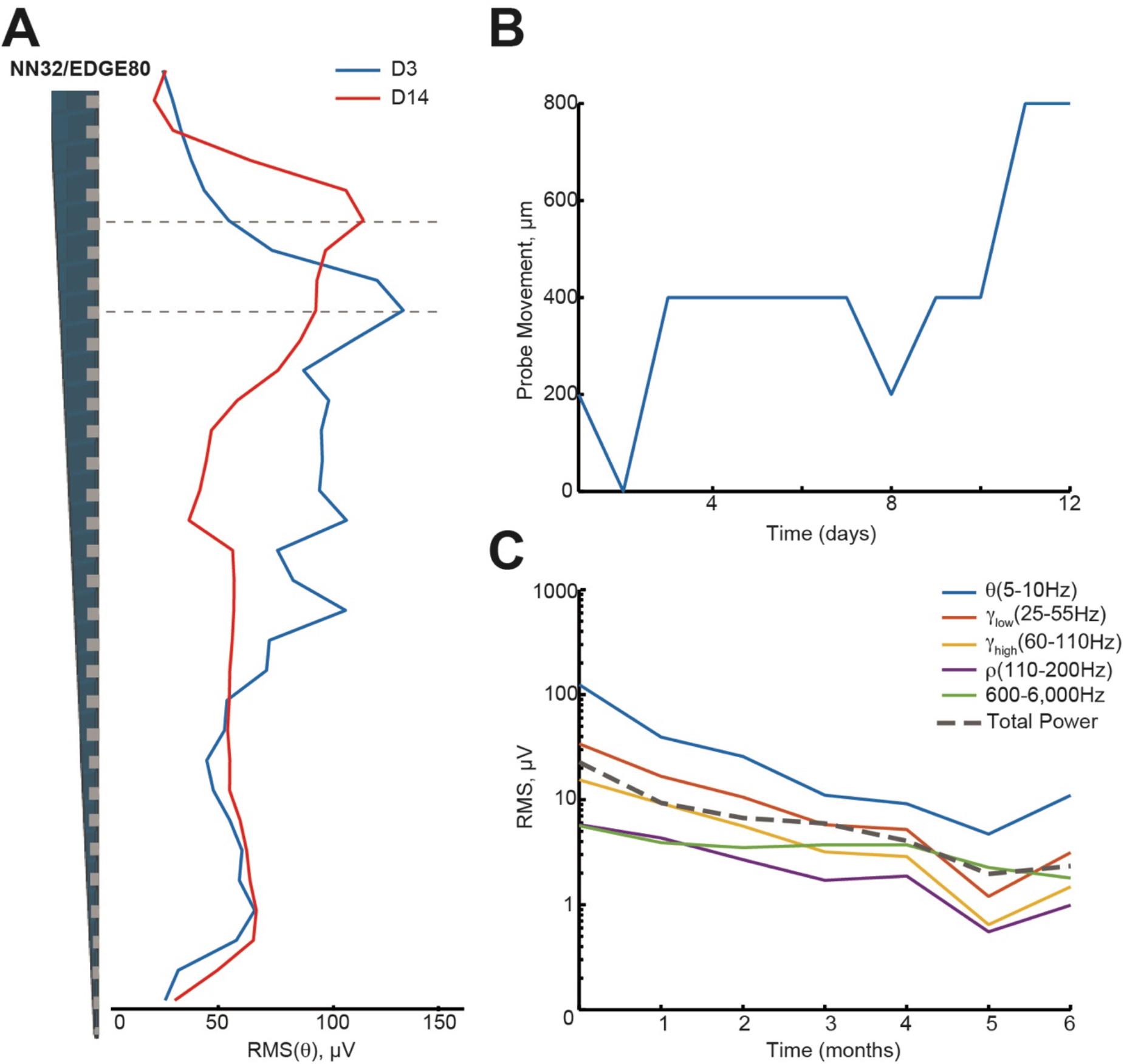
Hippocampal oscillations power decreases over months following chronical implantation. **A)** Over a period of two weeks, a chronically implanted NN32/EDGE80 silicon probe moved deeper into the dorsal hippocampus. One day post-implant, theta oscillation (~ 4 – 10 Hz) was maximal at channel 8 (blue trace). Over the course of two weeks, the theta peak moved up 3 electrode sites (red trace). A dotted line indicates the pyramidal CA1 layer (gray). **B)** The silicon probe’s drift was calculated for NN/EDGE80 silicon probe as a drift of theta power, which peaks in stratum-radiatum (blue trace). NN/EDGE80 probe moved down into the dorsal hippocampus for 600 μm in a course of 12 days following chronic implantation surgery. **C)** Overall power of hippocampal oscillations (θ (5 – 10 Hz), γ_low_ (25 – 55 Hz), γ_high_ (60 – 110 Hz), ρ (110 – 200 Hz), and 600 – 6,000 Hz) in dorsal hippocampus decreased over months, with significant drop in the first month. The total power is shown with a dotted line.

#### Stability of Single Units over Time

Next, we evaluated the amplitude and number of single units recorded from the porcine hippocampus over time (Figure 4). Since the 600 – 6,000 Hz power band representing unit activity decreased over time (Figure 3C), we compared the ability of various silicon probes to detect single unit activity during the first month post chronic implantation. Off-line spike detection and sorting was performed on the wideband signals (see Materials and Methods for details), with an example of raw signal (unfiltered, 1 – 15,000 Hz) recorded with NN32/EDGE80 silicon probes shown in Figure S2A. In planar silicon probes (ATLAS32/TET and NN32/TET), where the individual electrode sites were placed on a face of the designs, there were no single units recorded after a couple of days post chronic implantation. In an attempt to decrease the damage near electrode sites and increase the exposure of the electrode sites to the parenchyma, we designed NN32/EDGE style probe (Figure 1A). The individual electrode sites are strategically positioned on the edge of the substrate, potentially reducing the interference of the insulator with the surrounding signals [34].

**Figure 4.**
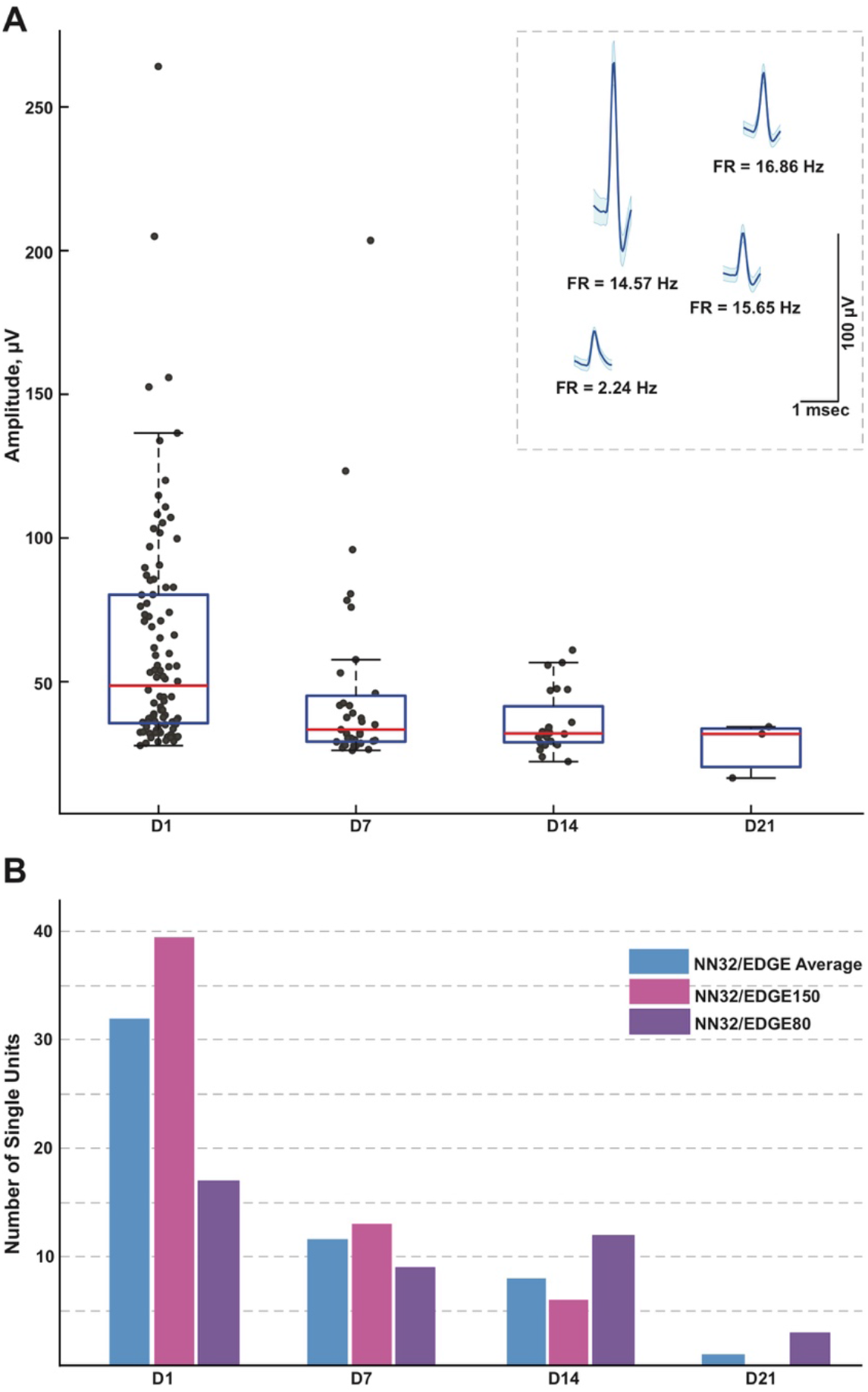
Single units recorded for three weeks post implantation. **A)** Single units were recorded in awake behaving pigs for up to three weeks post implantation (n = 3). While average amplitude of single units recorded with NN32/EDGE silicon probes at D1 post implantation was 63 ± 4 μV (mean ± SEM, range 27 – 264 μV, n_single units_ = 96), it decreased over time to 47 ± 6 μV (mean ± SEM, range 26 – 204 μV, n_single units_ = 35) at D7 and 36 ± 2 μV (mean ± SEM, range 22 – 61 μV, n_single units_ = 29) at D14. In addition, at D21 post implantation, single units were detected only from the narrower design silicon probe (NN32/EDGE80). The average amplitude of single units was 28 ± 6 μV (mean ± SEM, range 16 – 34 μV, n_single units_ = 3), just above a threshold for spike detection. Mean waveform and firing rates of representative single units are shown at D1 (inset). **B)** An ability to detect single unit activity was compared for NN32/EDGE150 and NN32/EDGE80 over three weeks period. NN32/EDGE150 silicon probes (n_single units_ = 40, pink, n = 2) had significantly more single units immediately after implantation (D1) than NN32/EDGE80 (n_single units_ = 17, n = 1, purple). However, there were no units recorded with NN32/EDGE150 after three weeks (D21), while NN32/EDGE80 silicon probes still detected single units (n_single units_ = 3, n = 1, purple). Total number of units averaged over time for combined NN32/EDGE style probes is also shown (n = 3, blue).

With NN32/EDGE silicon probes, single units were recorded for up to three weeks post implantation (Figure 4A). While average amplitude of single units recorded with NN32/EDGE silicon probes at D1 post implantation was 63 ± 4 μV (mean ± SEM, range 27 – 264 μ V, n_single units_ = 96), it decreased over time to 47 ± 6 μ V (mean ± SEM, range 26 – 204 μV, n_single units_ = 35) at D7 and 36 ± 2 μV (mean ± SEM, range 22 – 61 μV, n_single units_ = 29) at D14. In addition, at D21 post implantation, single units were detected only from the narrower design silicon probe (NN32/EDGE80). The average amplitude of single units was 28 ± 6 μV (mean ± SEM, range 16 – 34 μV, n_single units_ = 3), just above a threshold for spike detection (Figure 4A). The number of single units recorded with NN32/EDGE style silicon probes also decreased over the period of the first three weeks, with 64 % of single units disappearing by D7 and additional 31 % falling below detection threshold by D14 (Figure 4B). Immediately after implantation (D1), the 150 μm wide EDGE-style silicon probe (NN32/EDGE150) had more single units than similar silicon probe of just half the width (NN32/EDGE80, width = 80 μm) (NN32/EDGE150: n_single units_ = 40 vs. NN32/EDGE80: n_single units_ 17). However, after three weeks (D21), no units were detectable with NN32/EDGE150 silicon probe, while NN32/EDGE80 still recorded single units (n_single units_ = 3), suggesting a greater amount of tissue damage produced by the silicon probe with a larger cross-sectional area (Figure 4B).

#### Chronic Tissue Response to Implantation of Silicon Probes

To test how crosssectional area of the silicon probe (measured at the top electrode site) affects stability of neurophysiological signals recorded chronically (over time), we histologically evaluated tissue damage in the porcine hippocampus produced by silicon probes of various designs (Figure 5). Microscopic examinations performed following acute silicon probe insertions (n = 3, multiple electrode types within 2 – 5 hours) demonstrated (as expected) a degree of tissue disruption with associated hemorrhage around the probe track (Figure 5A). In addition, axonal pathology could be identified immediately adjacent to the site of silicon probe placement as evidenced by the pathological accumulation of APP, likely secondary to axonal transection causing acute interruption of transported proteins (Figure 5B). While, direct comparisons of chronic histological outcomes of different probe types were not performed, preliminary histological evaluations reveal the nature of the pathological response to silicon probes in situ over time in swine. Specifically, at 1.5 months post-implantation (n=4, multiple electrode types), hypercellularity was observed surrounding the silicon probe track. Notably, in one case the choroid plexus could be visualized entering the alveus, presumably having been translocated during the insertion as has previously been visualized (see Figure 8 in [35]) (Figure 5C). Immunostaining revealed both reactive astrocytes, with increased immunoreactivity to GFAP, and IBA-1 positive cells with the morphological appearance of reactive microglia (D-F). Interestingly, Van Gieson’s staining also revealed collagen surrounding the silicon probe track (G). Notably, these features were also observed even at 6 months post-implantation, regardless of the type of silicon probe (n=2) (Figure 5, H-K).

**Figure 5.**
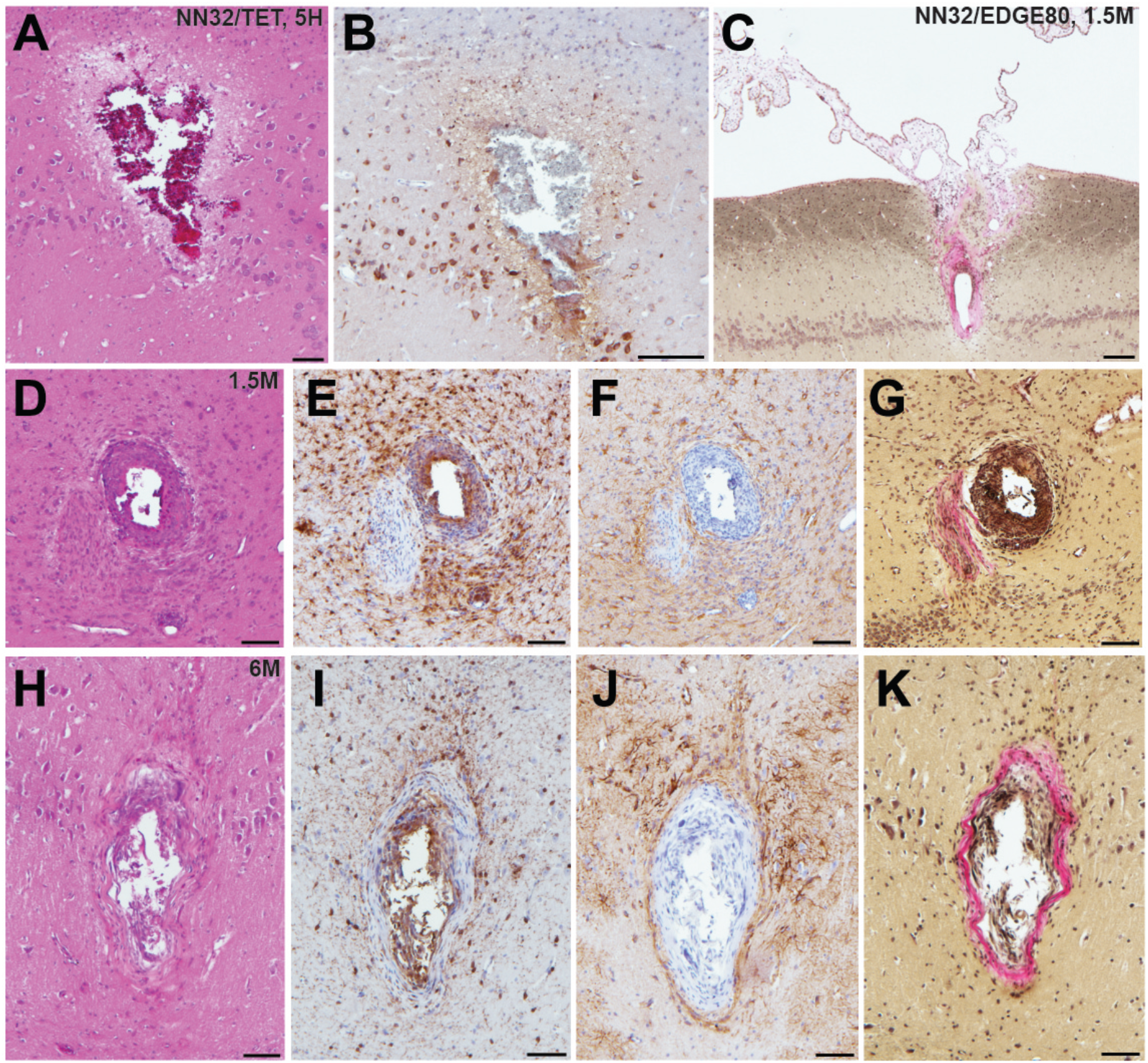
Acute and chronic tissue response to implantation of silicon probes in a large animal. Histopathological responses of multichannel silicon probes were microscopically examined at acute and chronic time points. **(A)** H&E staining showing silicon probe track with associated hemorrhage acutely (<5h) following insertion of NN32/TET electrode. **(B)** APP immunohistochemistry showing the same region as (A). Note the axonal swellings indicative of axonal transport interruption, likely secondary to axonal transection during insertion. Surrounding pyramidal neurons also show increased APP immunoreactivity. **(C)** Van Gieson stain showing silicon probe track 1.5 months following implantation of NN32/EDGE80 probe where collagen is pink. Notably, there appears to be choroid plexus abnormally located within the alveus. **(D-G)** Neuropathological findings in NN32/TET silicon probe track, 1.5 months following implantation, demonstrating **(D)** lesion with **(E)** IBA-1 positive cells displaying morphological features of activated microglia, **(F)** GFAP immunoreactive cells with features of reactive astrocytes and **(G)** Van Gieson staining demonstrating the presence of collagen (pink). **(H-K)** Silicon probe track 6 months following implantation of NN32/TET, again showing surrounding activation of both microglia **(I)** and astrocytes **(J)**, as well as the presence of collagen **(K)**. Scale bars: A, B, D – K: 100 μm; C: 200 μm.

### Novel Multichannel Silicon Probes Designed to Improve Outcome of Chronic Implantation in Large Animals

To increase unit yield and to improve longevity of neurophysiological signals (local field oscillation and single unit activity) over time and to reduce pathological tissue response to chronic silicon probe implantations, novel 64-channel flexible silicon probes were developed in a collaboration with SB Microsystems and Cambridge NeuroTech and tested acutely (under anesthesia) in pigs.

#### Lithography Process Defines Cross-Sectional Area of Chronic Silicon Probes

The lithography and metal etch or liftoff technologies available for a given silicon probe fabrication process determine the size and spacing of the conducting lines, and therefore the device cross sectional area for a given channel count. Based on our experience with chronic probe implants in large animals, probes with cross sectional areas on the order of 10^4^ square microns (as is the case for a 100 μm thick and 250 μm wide probe (Figure 1B)) are simply too damaging to tissue to reliably record single units on a chronic basis, and most practical recordings are carried out with probes that are 5 – 10 times slenderer than that. To our knowledge, past efforts at large animal probes have utilized contact lithography that limited interconnect pitch (width of a single conducting trace plus space) to 3-4 microns, leading to a practical upper limit of 32 channels per penetrating shank, given these constraints in cross sectional area. For example, 32-channel silicon probes designed by ATLAS Neuroengineering for large animal recordings (ATLAS32/TET) were 100 μM thick and made of silicon throughout the length of the probe (Figure 1B). While 32-channel silicon probes designed by NeuroNexus (NN32/TET, NN32/EDGE150 and NN32/EDGE80) were only 50 μM thick, they became brittle beyond 15 mm. This necessitated a novel interface cable solution whereby a high-density micro-cable was threaded through a 250 μm stainless steel tube and bonded to the contacts at the end of the probe, allowing the probes to be produced in lengths great enough to reach any part of the large animal brain (Figure 1B).

To reduce damage to brain tissue caused by insertion and chronic placement of silicon probes in the deep brain structures (hippocampus), we aimed to custom-design novel multichannel silicon probes with minimum cross-sectional area suitable for insertion. The goal was to reduce the profile of the probe, potentially reducing damage to the neural interface and inflammation. Increasing a probe’s thickness imparts greater mechanical stiffness of the silicon probe which benefits insertion, but causes a further compliance mismatch between probe and brain tissue, potentially contributing to gliosis and local cell death for chronic implementations. In order to test stiffness/insertion abilities empirically, mock versions of CAMB64 silicon probes were created to have a fixed width of 80 μm, while the thickness of the mock probes varied in a range of 25 – 100 μm. While 25 μm thick mock silicon probes could be inserted into cortex, they failed to advance to the hippocampal structure. In contrast, mock silicon probes with thickness of > 35 μm were capable of insertion into the hippocampus. Based on the insertion ability and acute neuropathology, the thickness of 35 μm was selected for production of initial research silicon probes, giving a minimum dimension for the probes as stiffness increases linearly with width.

To potentially reduce tissue damage even further, we aimed to design silicon probes to have a full-length silicon rather than a standard “silicon plus guide tube” design. By reducing the tissue impact during insertion, smaller, less brittle silicon probes may also increase the survival rate of neurons in close proximity to the travel path. In order to produce longer silicon probes without the need for the extension tube, novel research silicon probes were manufactured using projection (“stepper”) lithography rather than contact lithography, allowing us produce 0.5 μm resolution features and changing the width / channel count tradeoff in our favor. However, the use of projection lithography for these large devices (up to 40 mm in length) meant that the 5X master reticle exceeded the maximum allowed size, and required us to develop a process for stitching multiple exposures of multiple reticles across the process wafer. This novel lithography process allowed for the full length of the probe to be manufactured in silicon. The mock version of these new probes was capable of insertion into the dorsal hippocampus without support of metal guide tubes, which also resulted in less damage to cortical tissue en route to the hippocampus. The new lithography technology used to create novel research silicon probes also allowed for a doubling of the number of sites on the probe, leading to the first 64-channel silicon probes for large animals that are made exclusively from silicon wafer at lengths capable of reaching much of the large animal brain. In addition, the individual electrode sites on CAMB64/EDGE and CAMB64/POLY-2 silicon probes were composed of gold (Au) coated with an organic polymer, lowering their impedance and decreasing the noise during recordings in comparison to other silicon probes used (Figure S2).

#### Multichannel Silicon Probes Designed to Improve Single Unit Isolation

While a linear design is fit for a wide range of applications, isolation of single unit activity may be difficult with only a linear site arrangement. To utilize commercial spike sorting software available at the time (SpikeSort 3D, Neuralynx), individual sites on silicon probes with a linear design had to be artificially grouped into sets of four (tetrodes) in order for single units to be sorted. Previously, single units recorded with 32-channel silicon probes of the earlier designs could not be properly isolated with spike sorting software available at the time, partially due to potential overlap between putative units between the artificial tetrodes.

To address this issue, we initially custom-designed silicon probes (ATLAS32/TET and NN32/TET) to have most of the individual sites arranged in four sites placed close together (tetrodes), allowing for high-quality cell discrimination in hippocampal recordings (Figure 1A). Extra electrode sites were added in between groups of tetrodes in order to maintain the laminar analyses. In this configuration, four tetrodes were placed in pyramidal CA1 hippocampal layer (top part of the probe), while three tetrodes were placed in granular cell layer (bottom part of the probe). As tetrode-style silicon probes were created with the individual sites arranged closely to form tetrodes, more single units were isolated but crossover of the units onto neighboring tetrodes were still observed occasionally in the CA1 layer due to the large size and dendritic arbor of these neurons. As modern spike sorting software became available (Klusta and phy software packages), we designed silicon probes with a linear layout of the electrode sites, which also helped to resolve laminar structure of the porcine hippocampus (NN32/EDGE150 and NN32/EDGE80, Figure 1A).

The advent of laminar spike-sorting software removed the necessity for specific tetrodes to be recreated in the silicon probes, however probe geometry is still an important part of the spike-sorting process [28, 36]. We therefore designed novel research CAMB64 silicon probes with two arrangements of the individual electrode sites: a linear style CAMB64/EDGE probe (148 μm width) and a poly-2 style CAMB64/POLY-2 probe (154 μm width, electrode sites are off-set by 21 μm), with tip profile of both probes reduced to a minimum (Figure 1A, inset). To answer an open question whether vertically offset probe designs are necessary for true resolution of single units using modern sorting algorithms, we compared single-unit separation of a linear vs. a poly-2 CAMB64 silicon probes (Figure 6). Again, off-line spike detection and sorting was performed on the wideband signals (see Materials and Methods for details), with an example of raw signal (unfiltered, 1 – 15,000 Hz) recorded with CAMB64/POLY-2 silicon probe shown in Figure S2B. Decreasing spacing between individual electrode site to 100 μm on both CAMB64/EDGE and CAMB64/POLY-2 style probes helped to sort single units with more precision (Figure 6A). The blue unit on the linear (left) or poly-2 (right) style probes could have easily been classified with the red unit had the spacing not revealed the higher amplitude action potentials at the same time stamps. Moreover, the offset geometry of the poly-2 design also appears to better separate single cells compared to the linear design (Figure 6B). Many separated clusters from the CAMB64/EDGE recording contained multiple units, which were not well isolated from each other with current spike sorting methods. While some clusters from CAMB64/POLY-2 probe also contained multiple units, the proportion of multi-unit clusters was less when compared to the linear design (33 % and 54 % respectively).

**Figure 6.**
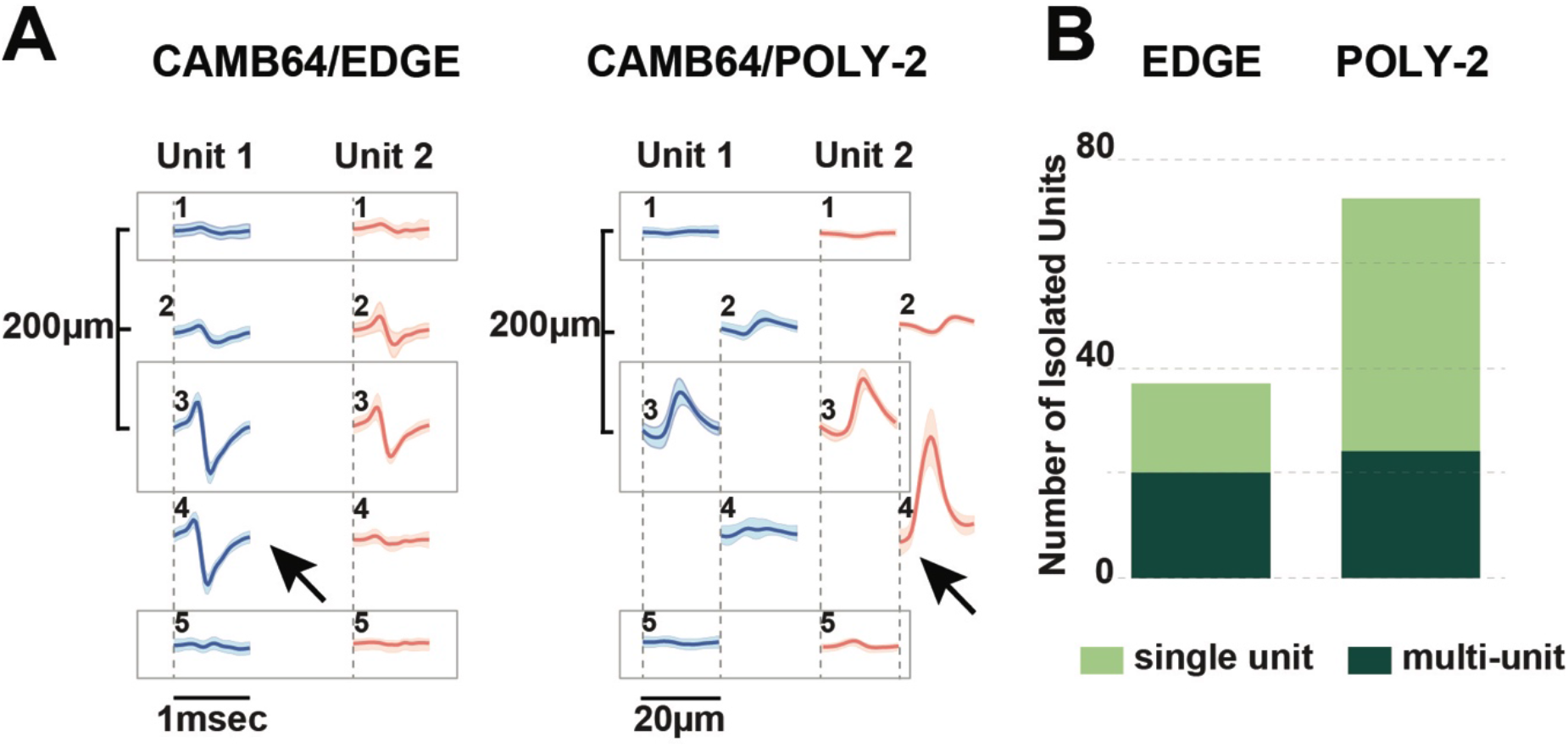
Neurophysiological signals recorded with novel research CAMB64 silicon from pig hippocampus. **A)** Single units were isolated for a linear and a poly-2 designs CAMB64 silicon probes. Decreasing spacing between individual electrode site to 100 μm on both CAMB64/EDGE and CAMB64/POLY-2 style probes helped to sort single units with more precision. The blue unit on the linear (left) or poly-2 (right) style probes could have easily been classified with the red unit had the spacing not revealed the higher amplitude action potentials at the same time stamps. **B)** Spike sorting on a recording with CAMB64/EDGE silicon probe yielded more multi-units (red, n = 20) than single units (green, n = 17). While some clusters on CAMB64/POLY-2 silicon probe contained multiple units (red, n = 24), two times more single units (green, n = 48) were identified. Also, the proportion of multi-unit clusters for poly-2 design was less when compared to the linear design (33 % and 54 % respectively).

## Discussion

Silicon multichannel probes designed and used for large animal hippocampal electrophysiology allow for a longer area of coverage in contrast to single site recordings, as well as revealing laminar structure and correlation of unit activity with field behavior. We have performed a retrospective analysis of chronically implanted pigs from our experiments in order to characterize the differences in electrophysiology over time with various probe sizes and geometries from different manufacturers. In addition, we have examined the neural interface out to 6 months post-implantation in order to assess whether neuropathology in the large animal hippocampus resembles that previously reported in rodents. Differing probe geometries were examined in a new higher density research probe in order to assess their ability for sorting units. Long-length silicon probes may also replace the need for multiple insertion acute experiments if the animal is chronically implanted and/or a drive is utilized.

Our initial attempts at chronic implantations using probes that were developed for acute work and prior to modern spike sorting methods for laminar probes were successful only for field recordings. Units that appeared during implantation surgery were either not present or were significantly attenuated even 24 hours post implantation. Histological examination suggested that the insertion damage to the neural interface, and sometimes even the cytoarchitecture, was significant with these larger dimension probes. In addition, attenuation of unit amplitude over time as well as field power in various frequency ranges is potentially explained by both cellular and collagen encapsulation of the probe as demonstrated at 6 weeks and 6 months post implantation. We therefore redesigned these probes to increase laminar coverage, as well as placing the electrodes on the edge of the probe in order to take full advantage of the contacts as has previously been described [34]. This change in dimension and geometry had the effect of increasing unit yields for the initial two to three weeks post implantation, with significant attenuation by the third week. Further reduction of the width by almost a factor of two at the top of active portion of the probe yielded a greater number of units at week three. It is an open question how much the placement of the edge electrodes increased unit longevity in relationship to the width change, but newer designs with offset spacing may allow for future examination of this relationship by comparing edge electrodes to those offset from the edge. Further detailed quantification of the differences in chronic neuropathology with various probe dimensions is also warranted to examine the potential correlation between these improved results and differences in the chronic neural interface response.

We also tested various designs of silicon probes available from and developed with the manufacturers for the ability to separate single units. Linear, tetrode plus linear and poly-2 site arrangement designs were evaluated for single unit vs. multi-unit cluster isolation. Spacing between individual electrode sites played a significant role as 200 μm spacing (as on NN32/EDGE style probes) increased the number of multi-units sorted. This could affect the precision of research studies focused on activity of hippocampal neurons, for example characterization of place cells or interneurons. Interestingly, some CA1 units spanned not two, but three electrodes, suggesting that large putative pyramidal cells that run parallel to the probe can be detected across greater distances than those previously reported, and further suggesting that lateral electrode separation for analysis in these structures may be helpful. Although most of the probes with a linear design site arrangement (NN32/EDGE150, NN32/EDGE80, and CAMB64/EDGE) were able to record neuronal activity of individual neurons, silicon probes with a tetrode (ATLAS32/TET and NN32/TET) and a poly-2 (CAMB64/POLY-2) design site arrangement had better cluster separation with currently available sorting algorithms, as the proportion of single units to multi-units recorded with a given probe increased. Previous acute examinations in large animals have also noted the usefulness of the parallel geometry in isolating units, even prior to the advent of new sorting techniques [22]. New sorting techniques, as well as drift associated with the semi-floating chronic preparation, suggest that parallel configurations may be optimal in order to maximize unit detection and separation in large animal laminar structures, and may be helpful as drives are developed for these probes.

Mock probe testing of the CAMB64 25 mm-long probes demonstrated that 35 μm shank thickness for 80 μm wide probes strikes an ideal compromise between ease of tissue penetration and appropriate targeting vs. the need to maintain small device dimensions. This allowed for production and testing of the 30 mm long 64-channel probes described above, which yielded substantially more units in the poly-2 style probe in acute testing. The balance between maximal stiffness, length and electrode density with minimal cross-sectional area in order to reduce damage to the neural interface remains a significant challenge in large animals. Designs such as the NeuroNexus’ NN32/EDGE probe and the new CAMB64 probe approach these challenges with different design solutions, but appear to have reached a critical threshold where the full depth can be reached with enough electrodes for sorting, and allowing for single unit detection out to a month post implant without being driven. Future chronic implantation of the CAMB64 probes will further test whether the trade-off of device width for increased channel count increases single unit yield, and for how long. These probes are also not at the minimal feature dimensions for this technique, as electrode yield in the initial testing run was also a consideration. Future dimensions of 80 μm wide, 64-channel probes may also increase yield by reducing damage to the neural interface, but this remains to be tested as well.

Passive laminar silicon probes (as described above) are but one solution to the problem of laminar recordings in large animals [37, 38]. Other proposed solutions that are either commercially available or being developed are the MicroFlex array and the Neuropixels probes, although the current Neuropixels probe is currently only 10 mm long and therefore cannot reach laminar deep brain structures in large animals such as the pig [39]. The main benefit of the MicroFlex array is a better biomechanical match to brain properties due to their flexible material, however the role of insertion trauma from electrodes or their carriers vs. chronic interface trauma due to mismatch in the modulus remains to be resolved. In addition, current reports in large chronically implanted animals have yielded fields, but not stable single units at present which remain to be demonstrated [40]. A future Neuropixels version for large animals would be of limited utility in its current form, as cell densities decrease in laminar structures as you move up on the phylogenetic tree [20]. In addition, it is not as practical to sample multiple structures along only one axis of a probe in the large animal as it is in rodents, necessitating multiple probes for multi-regional studies. Also, Neuropixels probes for chronic implants of a period of one year or longer may require different designs or materials to increase long-term biocompatibility [41]. Neuropixels probes cannot be used for translational research on neurological disorders that require neuromodulation as they are incapable of electrical micro stimulation, a technique also useful for probing the role of neural circuits in perception and cognition [42].

## Conclusions

We hope that this study will help to facilitate adoption of novel silicon multi-channel probes suitable for chronic implantations in large animals, by comparing silicon probes available for use in large animal electrophysiology, as well as comparing them to a new design and process. NeuroNexus EDGE style probes (NN32/EDGE80) were determined to yield the largest number of units of the available probes for acute and chronic recordings from laminar structures due a linear edge site arrangement, 6mm of coverage, and potentially reducing damage to the neural interface upon insertion. In addition, cross-sectional area was found to be one determinant of silicon probes’ performance. Novel CAMB64 silicon probes with a poly-2 design were found to have an even better single unit separation and a denser sampling of the laminar structure than existing linear probes. By increasing channel density, we were able to better visualize laminar structure and create offset geometries that enabled better unit sorting. Channel density, site arrangement, and physical profile of the silicon probe are all important factors to consider when designing probes for acute and chronic implantations to study laminar structures over time in awake behaving animals. We hope these results will lower the threshold for adoption in chronic implantations, by demonstrating consistent yields in laminar electrophysiological recordings in large animals using commercially available probes. We hope that in combination with new wireless technologies allowing for freely moving behavior, this will support new discoveries in both hippocampal and neocortical neurophysiology. In addition, better detection and understanding of laminar circuitry and changes in human disease (i.e. epilepsy, traumatic brain injury) are needed, and therefore viable electrodes need to be tested first in translational large animal models prior to clinical use.

## Author Contributions

Conceptualization, A.U. and J.W.; Methodology, A.U., P.K., T.H., B.J., and J.W.; Software, A.U., C.A., P.K, and J.W.; Validation, A.U., C.A., P.K, H.I.C., V.J., J.W.; Formal Analysis, A.U., C.A., C.C., K.G.; Investigation, A.U., C.A., P.K., K.G., J.W.; Resources, D.K.C. and J.W.; Data Curation, A.U.; Writing-Original Draft Preparation, A.U.; Writing-Review & Editing, A.U., C.A., K.G. and J.W.; Visualization, A.U. and J.W.; Supervision, J.W.; Project Administration, J.W.; Funding Acquisition, J.W.

## Acknowledgments

Authors would like to thank Matthew Sergison, Andy Tekriwal, and Maura Weber for their help with the experimental design and execution.

---

**Figure S1.**
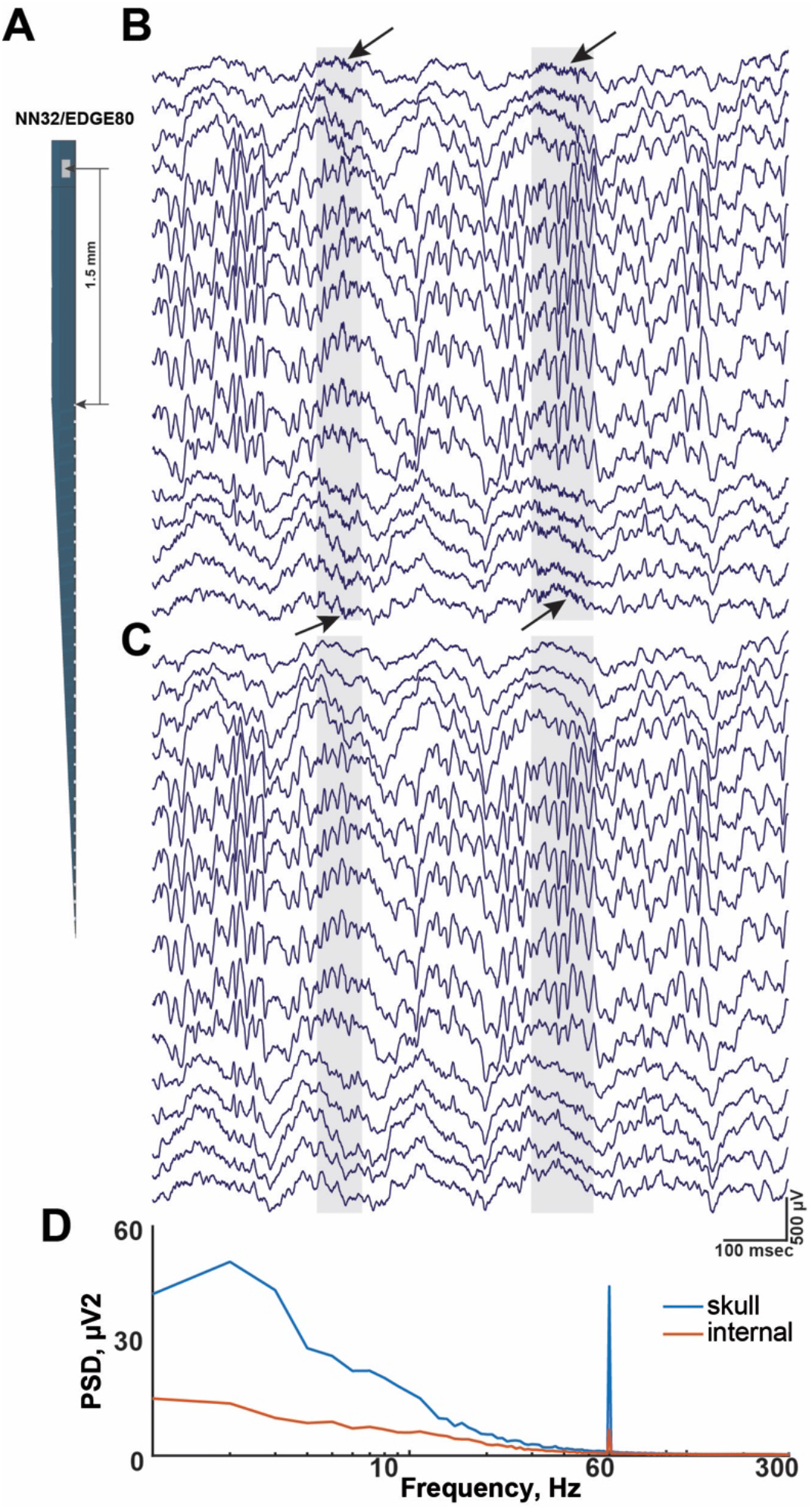
Internal reference on multichannel silicon probe reduces artifacts during chronic and acute recordings. **A)** All multichannel silicon probes used in the study were custom-designed to have a top electrode site substituted for a low-impedance reference site (Site Area = 4,200 μm^2^) and placed 1-2 mm above the most-proximal probe site. Schematic diagram of NN32/EDGE80 silicon probe showing location of internal reference. **B and C)** During awake behaving recordings in pigs, the internal reference eliminated movement-associated artifacts. The same 1 sec recording segment is shown referenced to a skull screw (B) vs. internal reference (C). **B)** The highlighted areas (gray) show “movement artifacts” detected on all 31 channels during awake recordings (arrows). Note that only 16 out of 31 channels are displayed. **C)** The noise associated with “movement artifacts” seen in B is eliminated after neurophysiological signals were re-referenced to the internal reference on NN32/EDGE80 silicon probe. **D)** Under anesthetized preparation, the internal reference on NN32/EDGE80 silicon probe eliminated most of the slow “drift” oscillations as well as 60 Hz frequencies peak, presumably from AC noise during acute recordings.

**Figure S2.**
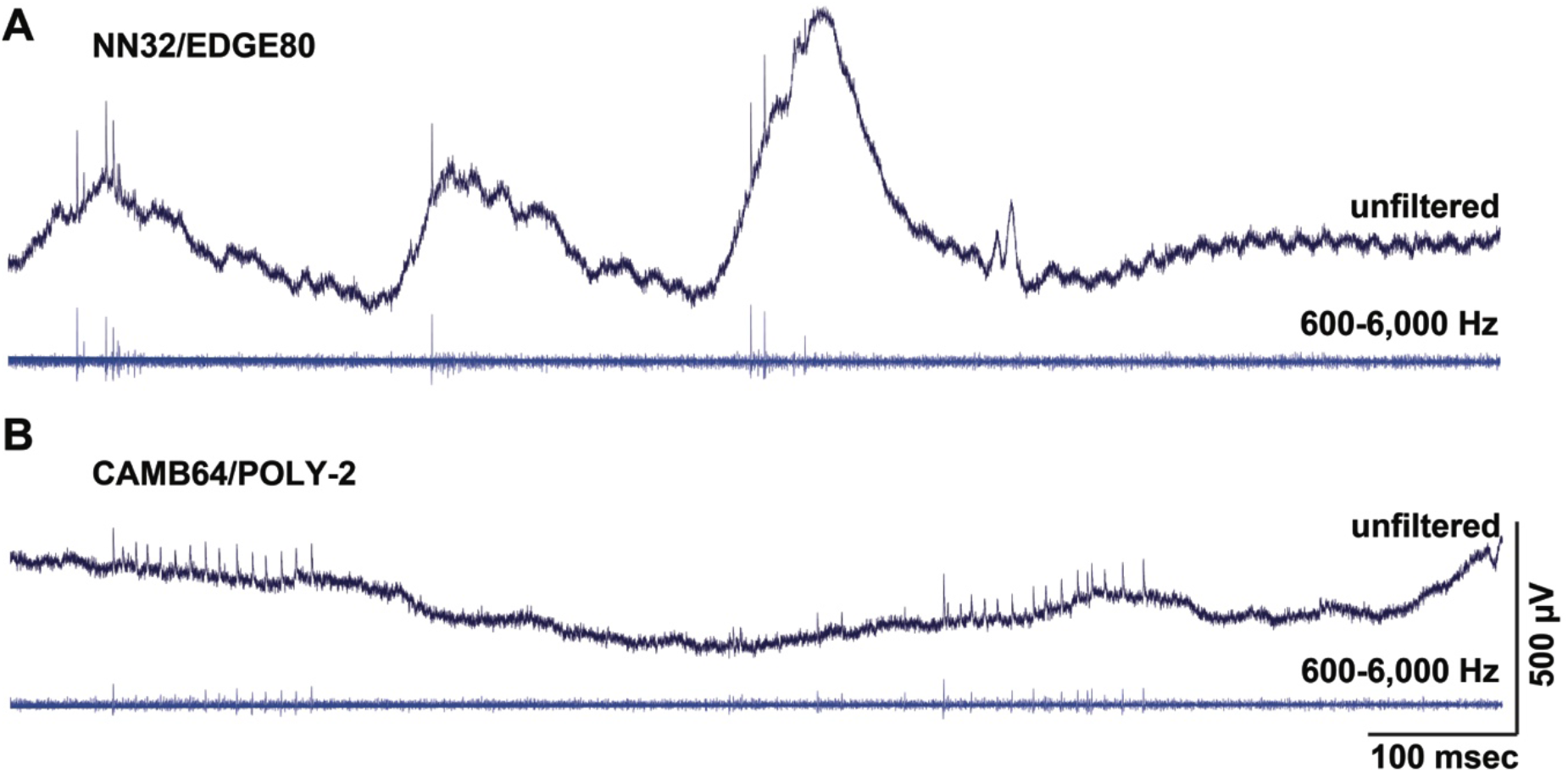
Multi-unit activity recorded with multichannel silicon probes designed for large animal electrophysiology. Recording profiles of multichannel silicon probes are displayed for NN32/EDGE80 and CAMB64/EDGE silicon probes. **A)** Representative traces recorded from a single electrode site on NN32/EDGE80 silicon probe that was located in the pyramidal CA1 layer show raw, unfiltered signal (0.1 – 9,000 Hz, top trace) and the filtered signal used to identify spiking activity (600 – 6,000 Hz, bottom trace). Multiunit and oscillatory activity can be seen on both traces. **B)** Representative traces recorded from a single electrode site on CAMB64/POLY-2 silicon probe that was located in the granular cell layer shows raw, unfiltered signal (0.1 – 9,000 Hz, top trace) and the filtered signal used to identify spiking activity (600 – 6,000 Hz, bottom trace).

